# Beyond kin killing: *Dickeya*-derived phage-tail-like bacteriocin P2D1 targets phylogenetically distant *Pseudomonas* spp

**DOI:** 10.1101/2025.10.07.680878

**Authors:** Marcin Borowicz, Jan Styn, Kacper Tomasik, Łukasz Rąbalski, Magdalena Narajczyk, Erwan Gueguen, Sylwia Jafra, Julie Baltenneck, Dorota M. Krzyżanowska, Robert Czajkowski

**Author notes:** These authors contributed equally to this work (Jan Styn, Kacper Tomasik).

## Abstract

Tailocins, phage-tail-derived bacteriocins, are increasingly recognized as potent mediators of microbial antagonism, yet their ecological scope beyond kin-targeting remains poorly understood. Here, we investigated whether P2D1, a tailocin produced by the plant pathogen *Dickeya dadantii* 3937, can act against environmental bacteria phylogenetically distant from *Dickeya* spp. Screening 480 soil and rhizosphere isolates from three distinct plant-associated habitats in Poland, we identified nine *Pseudomonas* spp. strains susceptible to tailocin P2D1. Whole-genome sequencing and phenotype profiling revealed that these isolates spanned multiple clades, including taxa related to *P. germanica*, *P. tensinigenes*, and *P. parakoreensis*. The *D. dadantii* mutant lacking genes encoding tailocin sheath and tube proteins lost antagonistic activity against *Pseudomonas* isolates, confirming that tailocins alone mediate the observed killing. Plant tissue assays revealed that six of the P2D1-susceptible strains were non-pathogenic and could mitigate *D. dadantii*–induced soft rot on potato. In contrast, three isolates related to *P. tensinigenes* were able to cause rot on their own under permissive conditions. Together, these findings demonstrate that P2D1 tailocin extends its activity to ecologically co-occurring but taxonomically distant *Pseudomonas*, suggesting that conserved receptors underline cross-genus targeting. More broadly, our results add to the limited evidence for tailocin activity beyond kin killing and therefore challenge the prevailing paradigm of kin-restricted tailocin specificity. They further suggest that tailocins may influence microbial community assembly across taxonomic boundaries, while their *in vivo* roles remain understudied.

## Introduction

Microbial communities in natural environments are shaped by intense competition for resources, with bacteria deploying a diverse array of strategies to outcompete their neighbors [1, 2]. Among these competitive factors, particularly intriguing yet poorly understood players are tailocins [3]. These phage tail-like bacteriocins are contractile nanomachines that, due to their structural similarities, are believed to be evolutionarily associated with bacteriophage tails [4]. Tailocins employ a single-hit killing mechanism facilitated by high-affinity recognition and membrane penetration in target bacterial cells, resulting in the rapid depolymerization of the susceptible cell and its ultimate death [5]. While tailocins production occurs across diverse bacterial species, their ecological role remains to be fully elucidated.

Tailocins are currently regarded as mediators of intraspecific competition, promoting kin killing through their specificity toward closely related strains, a context in which their activity has predominantly been characterized to date [6]. However, several studies have reported tailocins capable of acting across taxonomic boundaries [7, 8], suggesting their ecological role may be broader than previously assumed. This shift in perspective is particularly relevant in densely populated, taxonomically diverse microbial environments, where microorganisms continuously compete for limited space and resources [9, 10].

Soft Rot *Pectobacteriaceae* (SRP), including *Dickeya* spp., are well-characterized plant-associated bacteria known to produce tailocins [11]. We recently described a novel tailocin, dickeyocin P2D1, produced by *D. dadantii* strain 3937. Genetic clusters, such as those that encode the P2D1 tailocin, have been found to be widely distributed across *Dickeya* spp. [11, 12]. This suggests that P2D1 may play a significant ecological role in these bacteria, potentially conferring a competitive advantage within infected plant tissues and in other niches from which these bacteria are commonly isolated, such as soil, the rhizosphere, and aquatic environments [13, 14]. To date, SRP tailocins, including P2D1, have primarily been examined in the context of intraspecific competition, while their capacity to target non-kin bacterial taxa has not yet been systematically explored.

Here, we hypothesize that P2D1 tailocins can target phylogenetically distant environmental bacteria outside SRP, extending their ecological function beyond intra-species competition. To demonstrate this, we independently screened for P2D1 susceptibility in a pool of environmental isolates and constructed deletion mutants of *D. dadantii* 3937 that lacked genes encoding core structural components of the P2D1 tailocin to test their interaction with these isolates. With this approach, we assessed for the first time the potential widespread ecological role of tailocins, specifically P2D1 produced by *Dickeya* spp. strains.

## Materials and Methods

### Bacterial strains and culture conditions

All strains used in this study are listed in Table 1. Unless otherwise specified, routine cultivation of strains was carried out at 28 °C in Trypticase Soy Broth (TSB; Oxoid) with agitation at 120 rpm, or on Trypticase Soy Agar plates (TSA; Oxoid). *Escherichia coli* strains were cultured on LB agar supplemented with diaminopimelic acid (DAP) for the MFDpir strain. Chloramphenicol or ampicillin were added as required to maintain plasmids (Table S1).

**Table 1.** Bacterial strains used in this study.

| Bacterial strain | Description | Source |
| --- | --- | --- |
| ***Escherichia coli* K12** |  |  |
| DH5α | *supE44 lacU169 (Φ80lacZΔ M15) hsdR17 (rK mK ) recA1 endA1 gyrA96 thi-1 relA1* | Laboratory collection |
| DH5α λpir | λpir phage lysogen of DH5α | Laboratory collection |
| MFD*pir* | *RP4-2-Tc::(Δ Mu1::aac(3)IV-Δ aphA-Δ nic35-Δ Mu2::zeo) ΔdapA::erm-pir) ΔrecA* | [29] |
| ***Dickeya dadantii* 3937** |  |  |
| *D. dadantii* 3937 | Wild type, a representative member of Soft Rot *Pectobacteriaceae*, produces P2D1 tailocins | Laboratory collection |
| Δ3810 | Δ*Dda3937_03810*, deficient in tailocin sheat; collection number (Lyon): D643 | This study |
| Δ3811 | Δ*Dda3937_03811*, deficient in tailocin tube; collection number (Lyon): D644 | This study |
| Δ3810-11 | Δ*Dda3937_03810* Δ*Dda3937_03811* (double mutant); collection number (Lyon): D646 | This study |
| Δ3810-11 [pSJG] | Double mutant Δ3810-11 carrying the complementation vector pSJG | This study |
| **Enviromental isolates** |  |  |
| MC6 | A set of strains isolated from raspberry bush rhizosphere in Białogóra, Poland, 2024 (54,81533°N, 17,96580°E); susceptible to P2D1 tailocins from *D. dadantii* 3937 | This study |
| MD6 |  |  |
| ME6 |  |  |
| MB7 |  |  |
| Tul1A2 | A set of strains isolated from tulip flower bed rhizosphere in Gdynia Poland, 2024, (54,50774° N, 18,54757° E); susceptible to P2D1 tailocins produced by *D. dadantii* 3937 | This study |
| Tul2A7 |  |  |
| G3-3 | A set of strains isolated from cornfield soil in Częstochowa, Poland, 2024 (50,95110° N, 19,22670° E); susceptible to P2D1 tailocins produced by *D. dadantii* 3937 | This study |
| G3-19 |  |  |
| G3-39 |  |  |
| **Other** |  |  |
| *Dickeya zeae* NCPPB 3532 | Member of the Soft Rot *Pectobacteriaceae*, sensitive to P2D1 talocins and used as an indicator strain for their detection | Laboratory collection |

### Isolation of environmental bacterial strains

To obtain environmental isolates for subsequent testing for tailocin sensitivity, soil and rhizosphere samples were collected from three locations in Poland: tulip flower bed soil in Białogóra (54.81533° N, 17.96580° E), raspberry bush rhizosphere in Gdynia (54.50774° N, 18.54757° E), and cornfield soil in Częstochowa (50.95110° N, 19.22670° E). Samples used for microbial isolation were collected from environments with a potential presence of SRP bacteria [14]. Two grams of each sample were suspended in 4 mL of ¼-strength Ringer’s buffer (BioMaxima), shaken at 140 rpm for 30 min at room temperature, diluted 100× in the same buffer, and plated on 10% TSB agar. After incubation for 48 h at 28 °C, morphologically distinct colonies were collected, and the resulting 480 bacterial isolates were subcultured onto TSB agar plates under the same conditions to obtain pure cultures [15].

### Purification of P2D1 tailocins

P2D1 tailocin particles from *D. dadantii* 3937 were purified from mitomycin C-treated cultures using a previously described protocol [12]. The purified preparations were stored at 4 °C until use.

### Modified spot test for high-throughput P2D1 tailocin susceptibility screening

A fast and robust screening method involving 48-well plates (Greiner) was designed to test the new bacterial isolates for sensitivity to P2D1 tailocins. Each well was first filled with 20 μL of TSB medium. Next, a single bacterial colony of each tested strain was picked from a solid medium using a wooden toothpick and suspended evenly in the TSB droplet within the well. After inoculating all wells, 500 µL of molten soft-top agar (containing 30 g TSB and 7 g bacteriological agar (Oxoid) per 1 L), cooled to approx. 45 °C was added to each well. To facilitate immediate mixing of the inoculum with the freshly added medium, the multi-well plate was continuously agitated on a mini orbital shaker (Mini-Shaker, Biosan) (150 rpm) throughout the addition of the soft top agar. Once the agar had solidified, 2 μL of the purified P2D1 tailocins was spotted at the center of each well. Following 24 hours of incubation at 28 °C, the wells were inspected for zones of growth inhibition (clearance) indicative of tailocin activity. Wells inoculated with a P2D1-susceptible strain IFB 0117 [11] served as the positive control, while the negative control comprised the resistant P2D1 producer strain *D. dadantii* 3937. Isolates identified as sensitive during the screening were further tested using a classical spot assay [7].

### Genomic sequencing

Whole-genome sequencing of P2D1-susceptible isolates was performed on a MinION platform (Oxford Nanopore Technologies) using the wf-bacterial-genomes Nextflow workflow (v1.4.1). Genomic DNA was extracted with the Wizard Genomic DNA Purification Kit (Promega). Sequencing libraries were prepared according to the Oxford Nanopore ligation sequencing protocol and loaded onto R9.4.1 flow cells. Basecalling was performed with Guppy (v6.x, Oxford Nanopore). Reads were assembled *de novo* using Flye (v2.9.5) and polished with Medaka (v2.0.0). Assemblies were annotated with the NCBI Prokaryotic Genome Annotation Pipeline (PGAP, February 2025 release).

### Bioinformatic analyses

#### Taxonomic identification

Genome-based taxonomic identification and phylogenetic placement of the P2D1-sensitive isolates were performed using the JSpeciesWS web server [16] with FASTA genome sequences as input. First, the most closely related type strains in the GenomeDB database were identified for each isolate based on the highest Z-score correlation coefficient in pairwise tetra-nucleotide correlation (Tetra) analysis. Next, 5 type strains showing the highest Z-scores in Tetra-nucleotide analysis were subjected to pairwise Average Nucleotide Identity (ANI) calculations against the genomes of the strains under investigation. Both BLAST-based (ANIb) and MUMmer-based (ANIm) algorithms were applied. Additionally, the phylogenetic placement of the strains within the target genus was assessed based on 16S rRNA gene sequences (details are provided in Supplementary Dataset 1).

#### Dendrograms

To investigate the phylogenetic relationships between bacterial isolates, dendrograms were generated based on two independent datasets: genomic similarity and phenotypic profiles. Pairwise ANIm values (expressed as percent identity) were used to quantify genomic relatedness, and hierarchical clustering (agglomerative approach) was applied to group the strains accordingly. The same clustering method was used to analyze phenotypic data obtained from BIOLOG plate-based assays (described below), where positive and negative reactions were encoded as binary values (1 or 0). All clustering and dendrogram visualizations were performed using scikit-learn, SciPy, pandas, matplotlib, and numpy libraries [17, 18]. Python scripts supporting the analysis were developed with assistance from ChatGPT-4o (OpenAI) (Supplementary Script 1).

#### Comparative genomic analysis

To assess the genetic similarity among the nine P2D1-susceptible strains, we compared their genomes with respect to shared, accessory, and unique genes. The analysis was performed using the Pan-genome Explorer platform (https://panexplorer.southgreen.fr/cgi-bin/home.cgi) [19]. To assess the distribution of core and accessory genes, the PanACoTA pipeline was employed. Gene clusters predicted to be present or absent in individual environmental genomes were extracted and visualized using Venn diagrams generated in PAST software [20]

### Strain profiling with BIOLOG phenotypic microarrays

The ability of bacterial strains to utilize different carbon sources and their tolerance to various chemical stressors were assessed using the GEN III MicroPlate™ (94 phenotypic traits, including carbon utilization, chemical sensitivity, and physiological properties) and EcoPlate™ (31 carbon-source utilization traits) (Biolog) [21]. Plates were inoculated according to the manufacturer’s protocol and incubated at 28 °C. At 24 and 48 hours post inoculation, absorbance in each well was measured at 595 nm in the Epoch 2 microplate reader (BioTek). Results were normalized to the negative control and averaged across three biological replicates. A positive result was defined as an absorbance value at least twice that of the negative control.

### Assessment of phenotypic traits in P2D1-sensitive bacterial isolates

Bacterial isolates susceptible to P2D1 were investigated for selected traits on agar-solidified media plates: colony morphology was assessed on TSA and M9 0.4% glucose (MP Biomedicals), pectinolytic activity was evaluated on Crystal Violet Pectate (CVP) [22], siderophore production was assessed using Chrome Azurol S (CAS) agar [23, 24], and pyoverdine production was evaluated in King’s B [25].

### Microscopic imaging

The morphology of tailocin particles purified from the cultures of the wild-type *D. dadantii* 3937 and its mutants was investigated by transmission electron microscopy (TEM). TEM imaging was performed as described earlier [26] using the Tecnai Spirit BioTWIN microscope (FEI).

### Construction of P2D1-deficient mutants

P2D1 production in *D. dadantii* 3937 was abolished by generating in-frame deletion mutants in the loci *Dda3937_03810* (alternative locus designation *DDA3937_RS12110*) and *Dda3937_03811* (*DDA3937_RS12115*). The two genes encode the tail sheath protein (ADM98779.1) and tail tube protein (ADM98780.1) of the tailocin, respectively [12]. Additionally, a double mutant deprived of both genes was constructed. The deletions were performed using the pRE112 suicide plasmid (Cm^R^) carrying the *sacB* gene to enable counter-selection[27] (Supplementary Table S1). Procedures were analogous to those described earlier for *D. solani* [27]. Two PCR fragments corresponding to the upstream and downstream 0.5-kbp DNA of the gene(s) to be deleted in *D. dadantii* 3937 were amplified using the Primestar master mix (Takara) and cloned into SacI/KpnI digested pRE112 using the TEDA method [28] (list of oligonucleotides available in Supplementary Table S2). Chemical ultracompetent DH5α λpir cells were prepared with the Mix & Go! *E. coli* Transformation Kit using standard procedures (Zymo Research). Transformants were selected onto an LB plate supplemented with chloramphenicol (20 µg mL^−1^) and screened for the presence of the target construct by colony PCR with primers L762/L763. Constructs were extracted, confirmed by restriction map and Sanger sequencing, and then transferred into the competent *E. coli* strain MFDpir [29] prepared with the TSS method [28]. *E. coli* MFDpir produces the RP4 conjugation machinery, which allows the transfer of the suicide plasmid into *D. dadantii 3937* by conjugation. For conjugation, colonies of *D. dadantii* 3937 and MFDpir were mixed in the same proportion in 500 µl LB and centrifuged for 2 min at 8000 rpm. The pellet was resuspended in 90 µl LB with 5 µl diaminopimelic acid at 57 mg mL^−1^ and deposited onto an LB agar plate. After an overnight incubation at 30°C, the bacteria were resuspended in 1 mL LB, diluted in a 10-fold series from 10^-1^ to 10^-4^, and spread onto LB agar supplemented with chloramphenicol at 4 µg mL^−1^ to select the first event of recombination. Transconjugants re-isolated on this medium were then spread onto LB agar without NaCl, supplemented with 5% sucrose, and incubated at 20°C for 2-3 days to allow for the second recombination event. Sucrose-resistant colonies were then patched on LB-Cm plates to check for plasmid loss and streaked onto LB agar plates. The successful in-frame deletions were verified by colony PCR on purified colonies.

### Engineering a complementation construct for the P2D1 mutation

Previous RNAseq experiments [30] indicate that transcription of the Dda3937_03810 gene initiates at position 2751779, which is 16 bases upstream of its start codon. Furthermore, a strong transcription termination signal is located at position 2753545 – 39 bases downstream of the Dda3937_03811 gene. These findings suggest that *Dda3937_03810* and *Dda3937_03811* function together as an operon. Based on this data, we constructed a complementation plasmid designed to co-express both *Dda3937_03810* and *Dda3937_03811*. We cloned these genes, along with the 200 bp region upstream of *Dda3937_03810,* into the low-copy mobilizable plasmid pEGL332. This arrangement ensures that the native promoter for *Dda3937_03810* is aligned in the same orientation as the *plac* promoter within pEGL332. As a result, in the complementation plasmid, *Dda3937_03810* and *Dda3937_03811* are transcribed under the control of both their native promoter and the plac promoter, which increases the likelihood of transcription of both genes. The PCR fragment containing *Dda3937_03810* and *Dda3937_03811* were amplified with oligonucleotide pairs L1802/L1803 and then cloned into HindIII-linearized pEGL332 by TEDA [28]. The hybrid plasmid, designated as pSJG, was verified by restriction mapping and DNA sequencing. Then, it was transferred to *D. dadantii 3937* strains by biparental mating using MFDpir cell as the donor strain.

### Phenotypic comparison between wild-type *D. dadantii* 3937 and mutant strains

Phenotypes of the *D. dadantii* 3937 tailocin-deficient mutants, with and without complementation (Table 1), were compared to the wild-type strain. The comparisons included tailocin production, pathogenicity on potato tubers, and metabolic profiles assessed using GEN III MicroPlates and EcoPlates (Biolog), as described above. In addition, growth rates of the strains were evaluated at 28 °C in TSB and in M9 medium supplemented with 0.4% glucose.

### Antibiosis assay

The inhibitory activity of *D. dadantii* 3937 against environmental *Pseudomonas* spp. strains and *vice versa* was evaluated in an *in vitro* antibiosis assay on TSA plates, following the protocol [31]. After overnight incubation at 28 °C, cultures were examined for the presence of inhibition zones. The assay was performed twice, with three technical replicates each time.

### Virulence assays on potato tubers

#### Whole-tuber injection assay

To prepare the bacterial inoculum, cells from an overnight culture in TSB were harvested by centrifugation (4200 RCF, 5 min) and resuspended in PBS buffer. The turbidity of the suspension was adjusted to 0.06 McF (ca. 2 x 10^6^LCFULmL^−1^). To prepare plant material, tubers cv. Gala were surface sterilized by immersion for 20 min in 5% commercial bleach (ACE, Procter and Gamble), followed by a double rinse in distilled water and air drying under laminar flow. Each tuber was inoculated by inserting a pipette tip containing 50 µl of the test suspension into the tuber (up to the level of liquid within the tip) [32]. Tubers inoculated with PBS buffer alone were used as a negative control. Ten tubers per treatment were used to assess the potential virulence of P2D1-sensitive environmental isolates and to compare the *D. dadantii* 3937 wild-type strain with its mutants. Inoculated tubers were placed in humid boxes (85 to 90% relative humidity). Samples were incubated at 28°C to enable the development of soft rot symptoms. After 72 h, the tubers were cut at the inoculation site, and the tissue macerated by bacteria, if present, was spooned out and weighed. The experiment was conducted twice under the same conditions.

#### Potato tuber slices assay

Virulence and biocontrol assay on potato tuber slices was performed according to a modified protocol from [33]. Surface-sterilized potato tubers cv. Gala were sliced, and three wells (5 mm in diameter) were created in each slice. The slices were then placed in large glass Petri dishes (18 cm in diameter) lined with Whatman 3 filter paper disks, cut to fit, and moistened with 5 ml of sterile distilled water. To prepare bacterial inocula, cells were collected from overnight TSB cultures (4,200 RCF for 5Lmin) and resuspended in 0.85% NaCl. Biocontrol activity was assessed by mixing equal volumes of pathogenic strain suspensions (adjusted to 0.03 McFarland, ≈10LLCFULmL⁻¹) and candidate biocontrol strains (3 McFarland, ≈10LLCFULmL⁻¹). For pathogenicity assessment and single-strain controls, treatments consisted of a single strain mixed 1:1 with sterile 0.85% NaCl. Each well was filled with 30 μL of the corresponding mixture, with nine wells inoculated per treatment using three slices from different tubers. The slices were incubated in a humid chamber at 28 °C for 48 hours, after which the diameter of tissue maceration around each well was measured. The experiment was performed twice.

## Results

### P2D1 tailocin-sensitive isolates were detected across all sampled locations

Out of 480 environmental isolates tested, originating from three locations, nine were susceptible to P2D1: four from raspberry bush rhizosphere soil (MC6, MD6, ME6, MB7), two from tulip flower bed soil (Tul1A2, Tul2A7), and three from cornfield soil (G3-3, G3-19, G3-39) (Fig. 1A).

**Figure 1.**
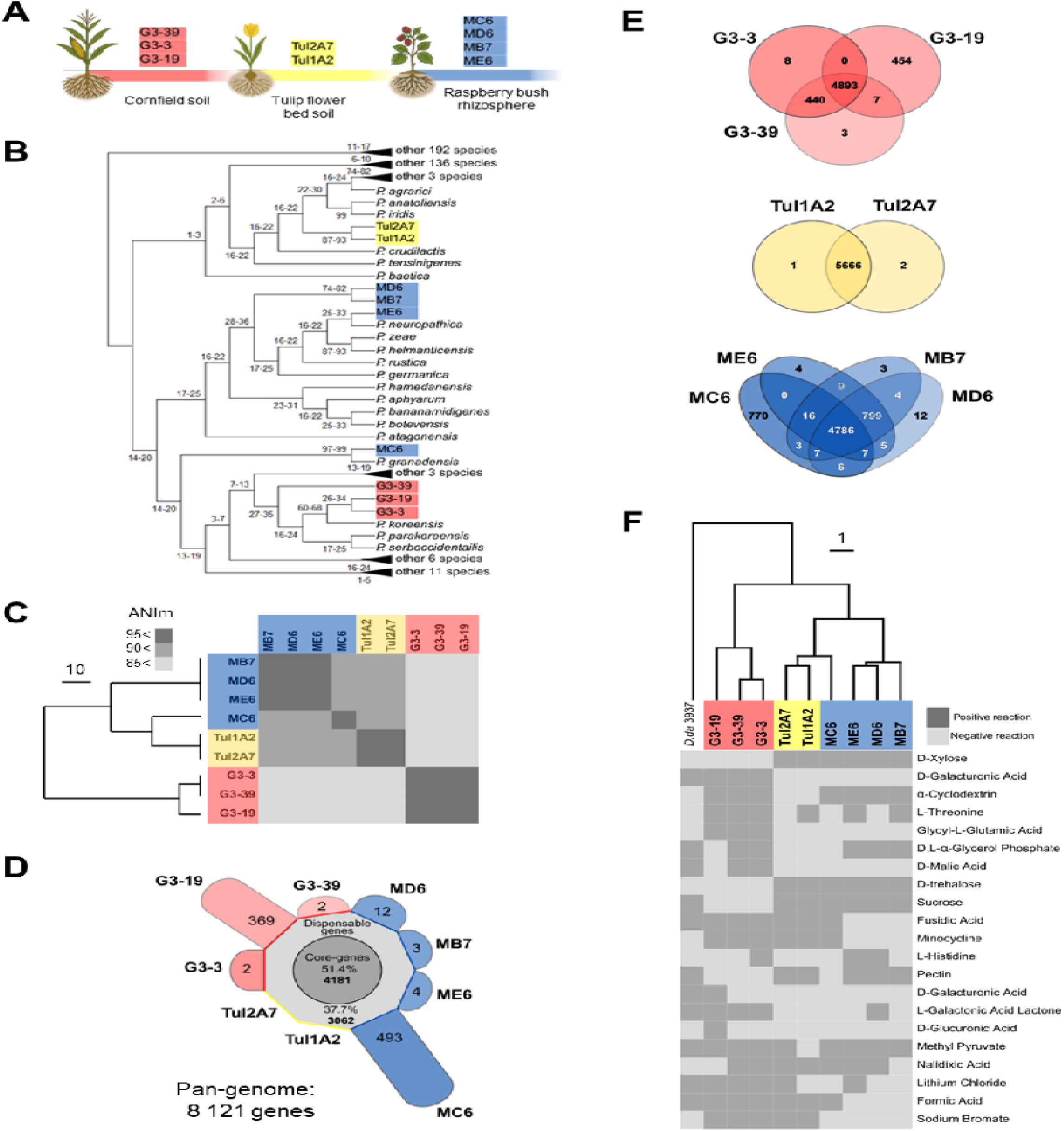
Genomic and phenotypic characterization of P2D1 tailocin-susceptible *Pseudomonas* isolates. **A.** Summary of isolation sources with the number of isolates obtained from each environment and the corresponding colour coding: red – cornfield soil; yellow – tulip flowed bed soil; blue – raspberry rhizosphere. **B.** Phylogenetic tree of *Pseudomonas* type strains based on 16S rRNA gene sequences. The tree was inferred using the Maximum Likelihood method with adaptive bootstrap support. For clarity, some branches are collapsed, while those containing the strains of interest are shown in full detail. The complete tree is available in Supplementary Dataset 1. **C.** Heatmap of pairwise ANIm values with hierarchical clustering. Higher values (darker shading) indicate greater genomic similarity, and the dendrogram illustrates the relationships among isolates. **D.** Distribution of core, accessory, and strain-specific (unique) genes in the pangenome. **E.** Venn diagrams showing: the shared gene clusters between isolates from distinct environmental origins; gene cluster overlap between strains from the same environment; and a comparison of gene content between a rhizosphere isolate from raspberry and isolates from tulip. **F.** Phenotype-based dendrogram derived from BIOLOG metabolic profiles (125 individual assays), illustrating phenotypic diversity among susceptible strains. For clarity, only traits showing variation across *Pseudomonas* spp. strains are displayed in the heatmap. *D. dadantii* 3937 was used as an outgroup. Complete results are provided in Supplementary Dataset 3.

### P2D1-sensitive isolates are members of *Pseudomonas* spp

Analysis of 16S rRNA gene sequences classified all nine P2D1-sensitive isolates within the genus *Pseudomonas* spp. (Fig. 1B; Supplementary Dataset S1). Whole-genome sequencing enabled higher-resolution taxonomic assignment based on ANI, the accepted standard for species delineation (species-level cutoff: ∼95–96%).

Three of the four raspberry rhizosphere isolates (MD6, ME6, MB7) were assigned to *P. germanica*, while the two tulip flower bed isolates (Tul1A2, Tul2A7) aligned with *P. tensinigenes* (ANI >95%) (Table 2). Strain MC6 also clustered closest to the *P. tensinigenes* type strain, but its ANIm (93%) and ANIb (92%) values fell below the species threshold. Cornfield soil isolates (G3-3, G3-19, G3-39) were most closely related to *P. parakoreensis*, yet similarly failed species assignment (ANIb 92%, ANIm 93%). Thus, while several isolates could be confidently assigned to known *Pseudomonas* species, others (MC6, G3-3, G3-19, G3-39) likely represent previously undescribed taxa. Several isolates: G3-3 and G3-39 from cornfield soil; MD6, ME6, and MB7 from raspberry; and Tul1A2 and Tul2A7 from tulip – exhibited ANIb values exceeding 99% with >99% alignment (Supplementary Dataset S2). This suggests that, although independently recovered from environmental samples, isolates within each source are in most cases clonal or near-clonal, with the notable exceptions of strains MC6 and G3-19.

**Table 2.**
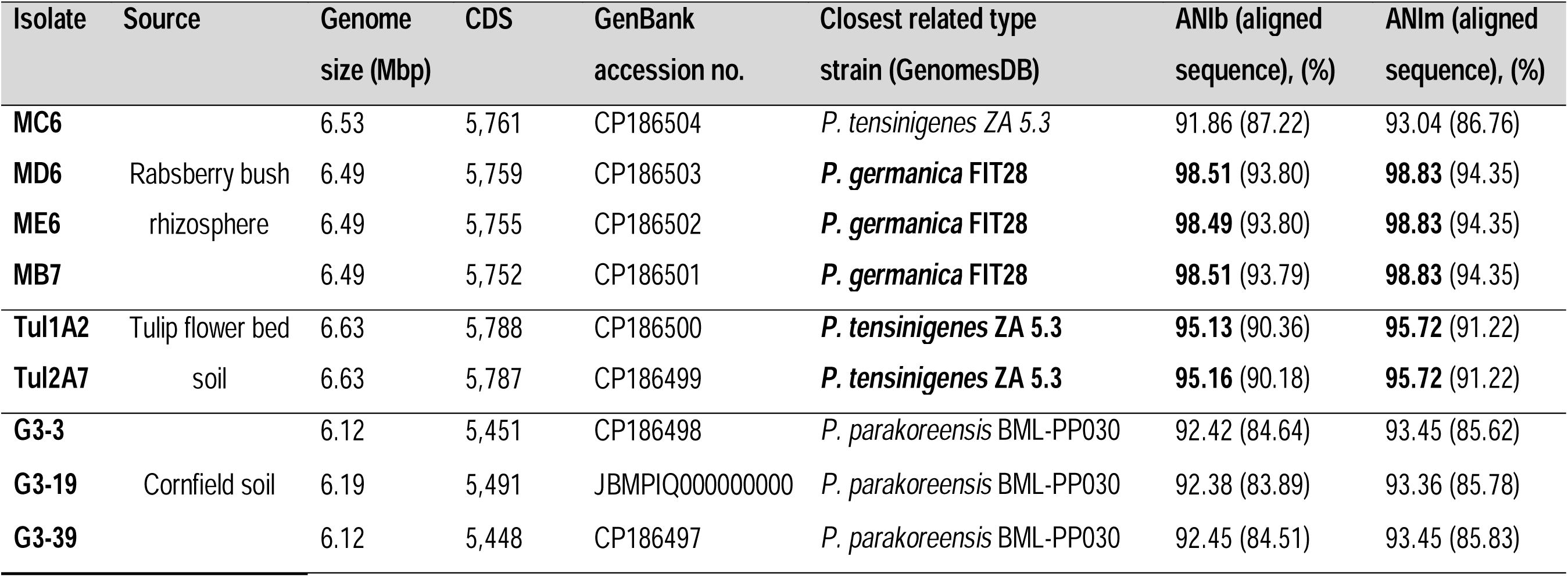
Genome characteristics and ANI values relative to the closest type strain.

Hierarchical clustering of ANIm values revealed three robust clades (Fig. 1C). Generally, isolates from a given sampling location clustered together, demonstrating phylogenetic consistency within habitats. An exception was strain MC6, which, although isolated from raspberry, clustered with the two tulip flower bed isolates. Despite this close clustering, ANIm values below 90% indicated that MC6 was conspecific with neither its environmental group nor the tulip isolates.

Within each environmental clade, the remaining strains exhibited ANI values greater than 95%, consistent with species-level similarity (Fig. 1C; Supplementary Dataset S2).

### P2D1-susceptible *Pseudomonas* spp. isolates form distinct groups based on genomic and phenotypic traits

Genome sizes of the analyzed *Pseudomonas* isolates ranged from 6.12 to 6.63 Mbp (Table 2). The collective gene pool of the nine strains comprised 8121 genes, including 4181 (51.4%) core genes shared by all strains and 3062 (37.7%) accessory genes present in at least two genomes (Fig. 1D). Strain-specific genes (10.9% of the total) were largely contributed by MC6 (493 genes) and G3-19 (369). In contrast, the tulip flower bed isolates carried zero, and the remaining five isolates harbored only 2–12 each (Fig. 1D). The gene count for G3-19 may be less reliable than for the other strains, as its genome is represented by contigs rather than a closed assembly.

Genomic diversity varied across clades: the tulip flower bed isolates were nearly identical, with only three unique genes in total, whereas the raspberry rhizosphere and cornfield soil clades displayed greater heterogeneity, each comprising two or more distinct P2D1-susceptible *Pseudomonas* strains (Fig. 1E). Out of 125 traits assayed across both BIOLOG plates, 21 showed variation among the isolates. Overall, strain-relatedness inferred from biochemical profiles was consistent with that inferred from genomic sequence analysis (Fig. 1F).

In plate-based growth assays, all nine isolates grew on TSA and on M9 with glucose as the sole carbon source. Colonies appeared glossy and mucoid, with abundant exopolysaccharide production most pronounced on TSA – a trait not universally observed among *Pseudomonas* spp. (Fig. S1A–B). All strains fluoresced on King’s B medium (Fig. S1C), consistent with pyoverdine production typical of fluorescent *Pseudomonas*. Only the three cornfield soil isolates (G3-3, G3-19, G3-39) grew and produced siderophores on CAS medium after 96 h, whereas the remaining six isolates showed no growth (Fig. S1D). No pectinolytic activity was detected on CVP medium at 24 h. Still, cavities appeared after 96 h in the tulip isolates Tul1A2 and Tul2A7 and in the raspberry isolate MC6 – the three strains that also cluster together in both genomic and biochemical analyses (Fig. S1E–F). These findings indicate genomic and phenotypic heterogeneity among the P2D1-susceptible *Pseudomonas* spp. isolates.

### *D. dadantii* 3937 tailocins attach to *Pseudomonas* cells, leading to cell puncture

Using transmission electron microscopy (TEM), we observed that P2D1 tailocins from *D. dadantii* 3937 attach to the surfaces of susceptible *Pseudomonas* cells, similarly to their binding on the positive control strain *D. zeae* NCPPB 3532 (Fig. 2A–F). These results provide direct ultrastructural evidence that *D. dadantii* tailocins physically bind to and interact with environmental isolates not related to SRP.

**Figure 2.**
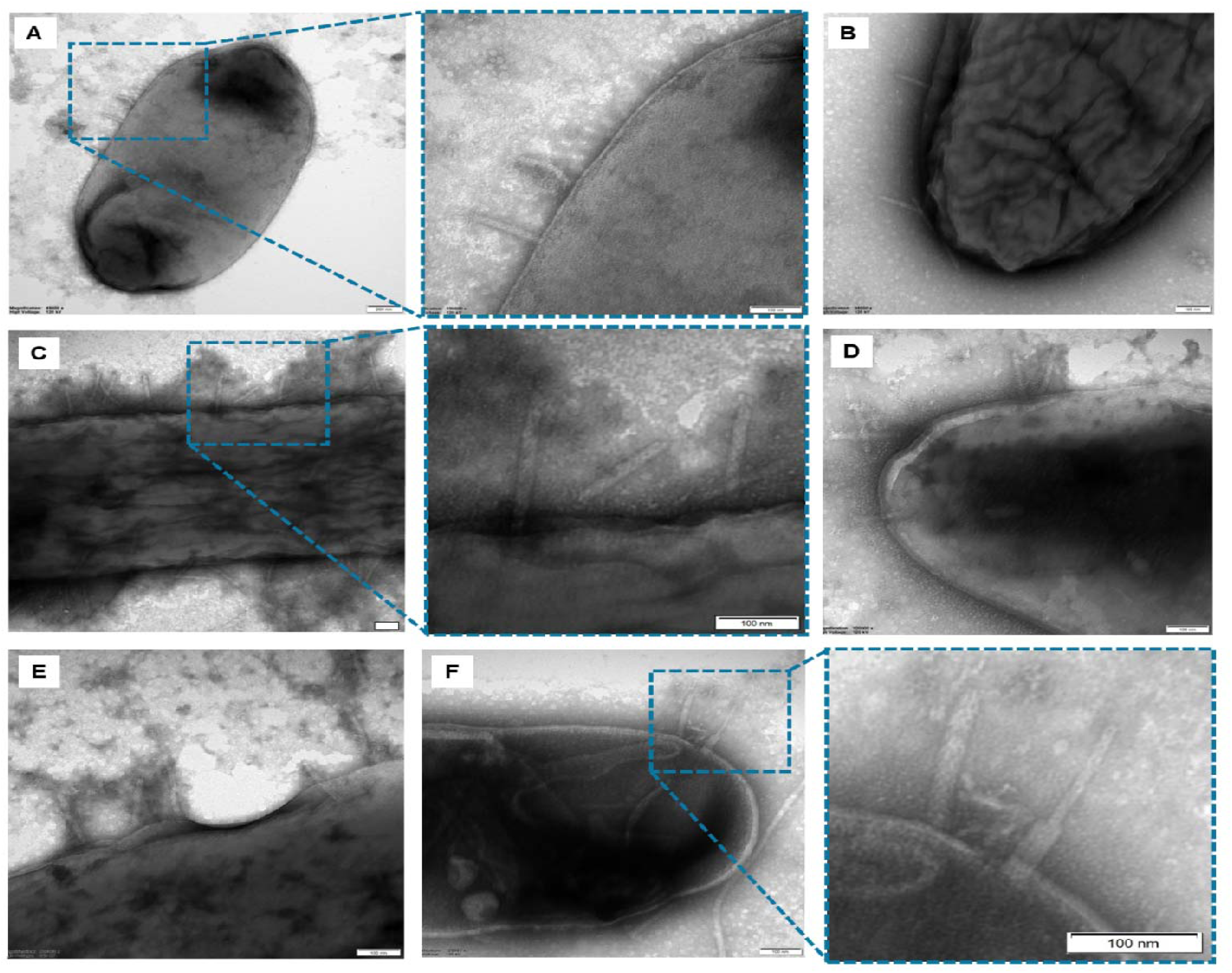
P2D1 tailocins of *D. dadantii* 3937 attached to the cells of susceptible environmental *Pseudomonas* spp. Panels: A - Tul1A2; B- G3-39; C - ME6; D, E - MD6; F - *D. zeae* NCPPB3532 (positive control). Scale bar: 100 nm.

### Tailocin-deficient *D. dadantii* mutants fail to kill environmental *Pseudomonas* spp.

To confirm that the inhibition of environmental strains by *D. dadantii* 3937 was mediated solely by tailocins, we constructed single (Δ3810, Δ3811) and double (Δ3810-11) mutants of *D. dadantii*, lacking genes encoding the sheath, the tube, or both genes, respectively (Table 1). None of these mutants produced complete (functional) tailocin particles (Fig. S2) or inhibited growth of the tailocin-susceptible SRP strain in soft-top agar overlay assays (Fig. 3). Aside from the absence of tailocin production, the mutants were indistinguishable from the wild type in growth rate (Fig. S3), virulence on potato tubers (Fig. S4, Fig. 4A), and biochemical profiles (Fig. S5). The double mutant (Δ3810-11) was selected for subsequent experiments.

**Figure 3.**
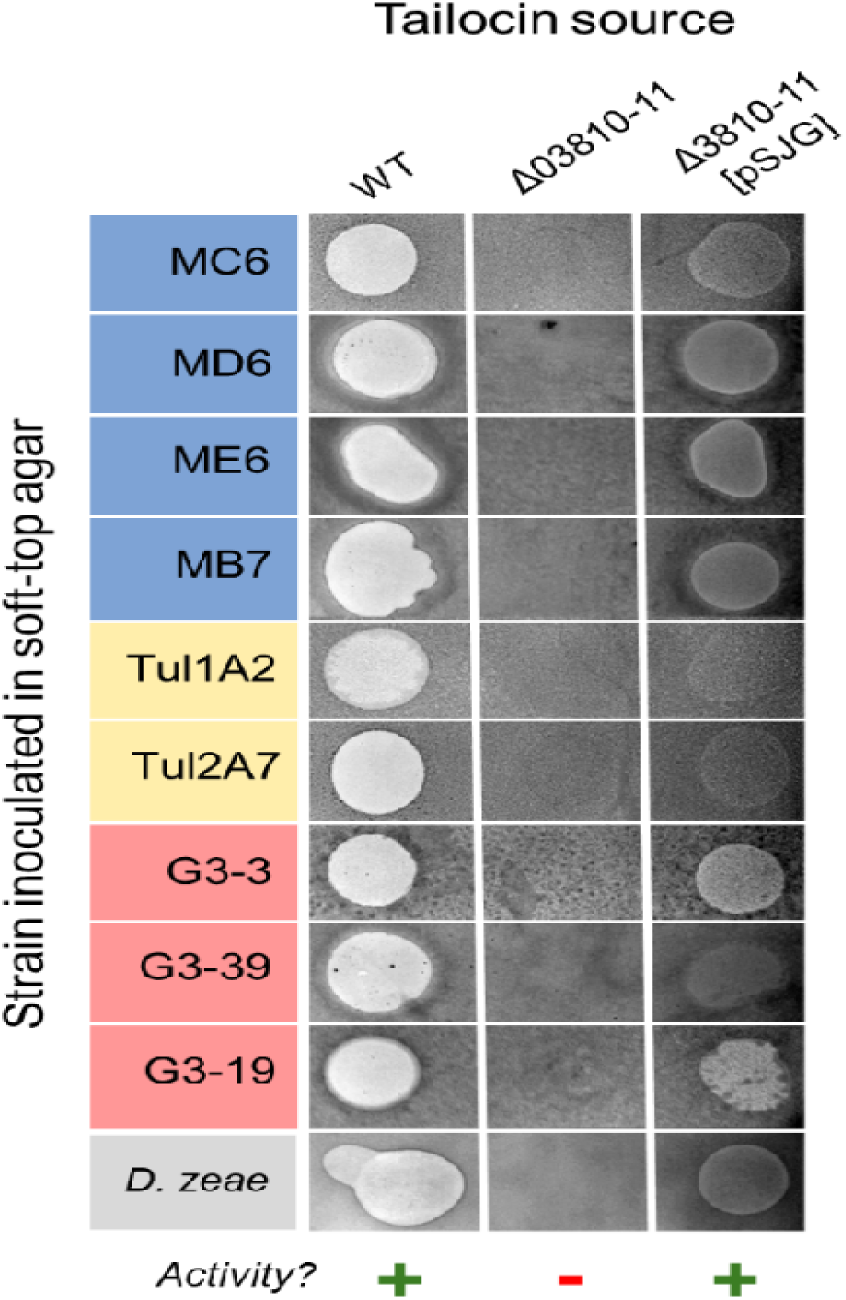
Activity of P2D1 tailocins isolated from *D. dadantii* 3937 WT, its P2D1- deficient mutant Δ3810-11, and a mutant with complementation plasmid pSJG. Tailocins were isolated from mitomycin C-induced cultures of the three tested strains and evaluated for activity against susceptible *Pseudomonas* isolates, as well as a control susceptible strain, *D. zeae* NCPPB 3532. The presence of a clearance zone indicates the presence of active tailocins in the tested preparation.

**Figure 4.**
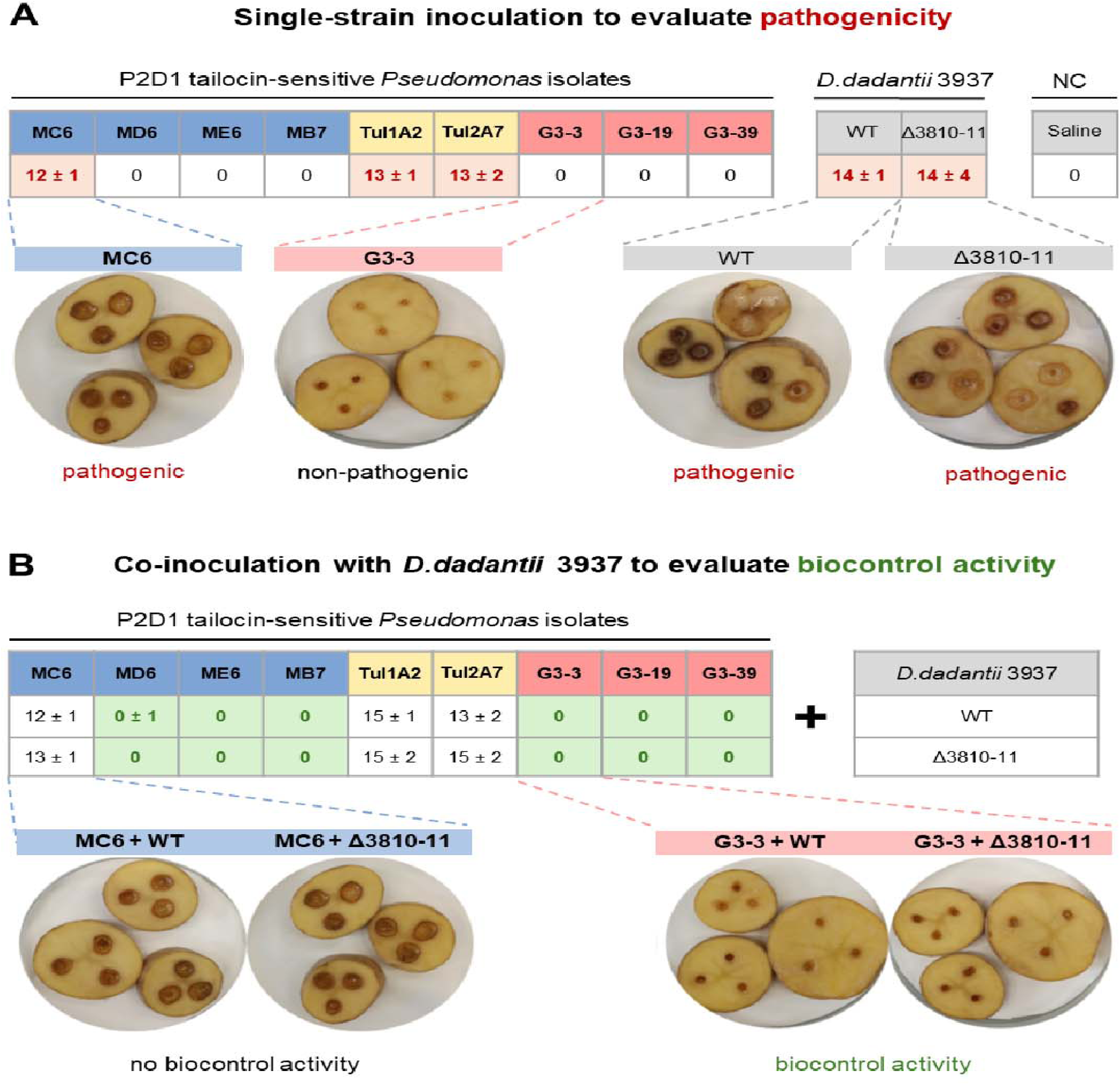
Effect of *Pseudomonas* isolates on potato tuber tissue in a slice assay. Panel A shows the ability of *Pseudomonas* isolates, *D. dadantii* 3937, and the 3937 tailocin mutant (Δ3810-11) to cause disease symptoms on tuber tissue when inoculated alone (pathogenicity). Panel B shows the potential of the *Pseudomonas* isolates to protect potato tissue from maceration when co-inoculated with the known pectinolytic pathogen *D. dadantii* 3937 (biocontrol activity). Median diameter of rotten tissue (mm; ± half interquartile range) is provided in the tables for all tested combinations. Alongside the tabulated results, the figure shows representative images depicting the characteristic appearance of healthy and diseased samples.

Tailocin preparations from induced Δ3810-11 cultures failed to inhibit any of the nine *Pseudomonas* isolates, in contrast to the wild type; however, complementation with plasmid pSJG (Table S2), which carries both disrupted genes under the operon’s native promoter, restored tailocin production and activity (Fig. 3, Fig. S2). Transformation with the complementation plasmid pSJG, most likely due to metabolic cost, resulted in a some unsignificant reduced growth rate of the transformed strains in M9 medium supplemented with 0.4% glucose, both in the mutant and in the wild-type host (Fig. S3).

Additionally, direct antibiosis was assessed using a plate assay to evaluate the production of antimicrobial metabolites. For both the wild type and Δ3810-11, only contact inhibition of *Pseudomonas* spp. isolates at colony borders was observed (Fig. S6A). These results indicate that, under the tested conditions, diffusible secondary metabolites of *Dickeya* do not contribute to antagonism against environmental *Pseudomonas*, supporting the conclusion that tailocins are the primary inhibitory factor. In a reciprocal setup, no inhibition of *Dickeya* strains by the tested *Pseudomonas* spp. was observed (Fig. S6B).

### *Pseudomonas* strains sensitive to tailocin P2D1 either cause potato decay or suppress *D. dadantii* in biocontrol assays

In the potato slice assay, three *Pseudomonas* spp. strains – Tul1A2, Tul2A7, and MC6 – caused tuber tissue maceration (Fig. 4A). These included two isolates originating from tulip flower bed soil and one raspberry isolate (MC6), all most closely related to *P. tensinigenes*. In the potato injection assay, which provides more microaerophilic conditions, only MC6 displayed mild pathogenicity within the experimental timeframe (72 h post-inoculation). In contrast, six isolates (MD6, ME6, MB7, G3-3, G3-19, and G3-39) not only lacked pathogenicity when inoculated alone (Fig. 4A), but also exhibited a biocontrol effect, significantly reducing soft rot symptoms caused by co-inoculated *D. dadantii* 3937 (Fig. 4B; with *Pseudomonas* isolates applied in excess, as typical for biocontrol assays). Together, these findings highlight the contrasting outcomes among P2D1-susceptible *Pseudomonas* isolates, ranging from independent pathogenicity to biocontrol of soft rot.

## Discussion

This study is the first to demonstrate that P2D1 tailocins produced by *the Dickeya* spp. can kill soil-associated *Pseudomonas* spp., which are phylogenetically distant from bacteria belonging to the Soft Rot *Pectobacteriaceae* (SRP) family. So far, tailocins have generally been described as possessing narrow host ranges, restricting their ecological role to competition with closely related strains [3, 34]. Only a few studies have demonstrated broader killing spectra, including *Pseudomonas fluorescens* tailocins suppressing *Xanthomonas vesicatoria* [8], a *Burkholderia cenocepacia* tailocin killing *Pseudomonas aeruginosa* [7], and *P. syringae* targeting *Erwinia amylovora*, *Xanthomonas perforans,* and the human pathogen *Salmonella enterica* [35]. Our present findings, derived from a study explicitly designed to search for such “off-target” effects, suggest that previously reported cases may not be rare exceptions but instead part of a broader and underexplored phenomenon with potentially significant ecological implications.

Among 480 environmental isolates tested, only nine (1.9%) were sensitive to P2D1 tailocins; yet these isolates were found across distinct environments and geographical locations. This so-called rare-but-widespread sensitivity pattern is consistent with results obtained in other studies, where tailocin-susceptible bacterial populations were typically low in frequency but widely distributed. For example, Yao and co-workers demonstrated that BceTMilo was active against 76 *Burkholderia* isolates, including both clinical and environmental samples, indicating its efficacy across multiple geographic and environmental contexts [7]. In other studies, maltocin P28 was active against 38 clinical and environmental *Stenotrophomonas maltophilia* strains [36], and tailocin from *P. syringae* USA011R targeted strains from distinct genera sourced from plant, clinical, and laboratory environments [35]. Likewise, R-type pyocins of *P. aeruginosa* targeted only a small fraction of clinical and environmental isolates; however, sensitive strains are found in different cystic fibrosis patients [37]. Furthermore, from an ecological perspective, finding susceptible isolates in geographically distant soils supports the idea that tailocins may act as selective forces in diverse plant microbiomes. Rhizosphere studies demonstrate that pyocins can significantly influence strain competition and community assembly, particularly under conditions of nutrient limitation [3].

Microscopic imaging revealed that P2D1 tailocins attach directly to the cell surface of susceptible *Pseudomonas* spp. isolates, resembling the interaction observed in the known susceptible strain *D. zeae* NCPPB 3532 [12]. This observation was supported by experiments with a P2D1-defective *D. dadantii* 3937 mutant, which failed to kill susceptible *Pseudomonas* spp. cells, thereby demonstrating that P2D1 tailocins alone are sufficient to mediate the observed killing. However, the specific surface determinants underlying P2D1 susceptibility in both *Dickeya* and *Pseudomonas* remain unidentified. Studies of *Pseudomonas* strains susceptible to tailocins produced by other *Pseudomonas* (pyocins) have shown that their binding specificity and target sensitivity are primarily governed by the structure, composition, and proper presentation of the lipopolysaccharide (LPS) O-antigen on the bacterial surface [38]. In the same model, conserved LPS chemotypes promote cross-species binding and susceptibility to tailocins among *Pseudomonas* strains [38]. In *Dickeya* spp. and the related *Pectobacterium* spp., LPS has been implicated in sensitivity to certain, but not all, bacteriophages [39, 40]. By contrast, the determinants of tailocin susceptibility in these bacteria remain uncharacterized. Identifying the surface features, particularly those shared between *Dickeya* spp. and *Pseudomonas* spp. that enable P2D1 cross-reactivity, may represent an important direction for future research, as described in other bacterial systems [41, 42].

In plant assays, P2D1-susceptible *Pseudomonas* spp. showed contrasting effects on plant tissue, both when inoculated alone and when co-inoculated with *D. dadantii* 3937. Three environmental isolates behaved as opportunistic soft-rot pathogens, whereas six others exhibited attenuation of *D. dadantii*-induced soft rot in potato tubers. These dynamics mirror the known dual roles of *Pseudomonas* spp.: some species (e.g., *P. marginalis*, *P. palleroniana*) cause soft rot [43, 44], whereas others serve as biocontrol and plant-beneficial agents [45]. Importantly, *Pseudomonas* spp. are known to co-occur with *Dickeya* spp. and *Pectobacterium* spp. in rotting potato tissues as part of a polymicrobial “spoilage microbiota” [46]. Several pectinolytic *P. fluorescens* strains have been identified alongside *Pectobacterium* spp. in diseased potato tubers in Kenya, confirming that both strains can coexist in rotten potato tissue[47]. At the same time, other *Pseudomonas* spp. are known biocontrol organisms: *P. chlororaphis* suppresses *Dickeya* spp. virulence by quenching quorum-sensing signals [48], while *P. fluorescens* eliminates competing *Pectobacterium* through a Type VI-secreted amidase, thereby protecting potato tubers [49]. Consistently, co-inoculation of antagonistic *P. fluorescens*, *P. putida* or *P. donghuensis* strains has been shown to significantly reduce potato blackleg/soft-rot severity caused by SRP pathogens [15, 31, 33, 50]

It is therefore evident that P2D1-susceptible *Pseudomonas* spp. isolates can compete with SRP, including *D. dadantii,* on different planes - either as alternative pathogens of potato tissue or as antagonists mitigating soft rot. Our results, therefore, indicate that by targeting *Pseudomonas* with divergent ecological functions, P2D1 tailocins could function as potent modulators of both microbial competition and plant health.

Taken together, our findings indicate that tailocins produced by plant-associated bacteria can act against ecologically co-occurring competitors across genera, rather than being restricted to genetically closely related strains. Clusters encoding P2D1-like tailocins are widespread across *Dickeya* spp., suggesting positive selection for this trait [11]. Such an antagonistic capacity may be particularly advantageous for soft-rot pathogens, which cycle between insect, aquatic, soil, and plant niches and repeatedly encounter diverse resident microbiota [13, 14, 51]. Likewise, tailocin production is inherently costly, as it requires cell lysis and can leave producing populations vulnerable to competition by insensitive community members in mixed habitats [52]. Consequently, the net benefit of tailocin deployment likely depends on spatial structure, timing, and the fraction of cells committing to production [53]. At the same time, the precise contribution of P2D1 to *Dickeya* success during niche invasion remains unresolved. Future *in vivo* studies will be crucial to determining the ecological relevance of tailocins and their role in shaping microbial community structure.

## Data availability

The *Pseudomonas* spp. genome sequences have been deposited in NCBI GenBank under accession numbers CP186497 - CP186504 and JBMPIQ000000000.

## Author Contributions

M.B. supervised laboratory experiments by undergraduate students, performed experimental work, carried out bioinformatic analyses, contributed to figure preparation, analyzed data, drafted the manuscript, and contributed to funding acquisition. J.S. and K.T. collected samples, performed the screening for tailocin susceptibility, and participated in laboratory experiments. Ł.R. performed DNA sequencing on the isolates. M.N. performed transmission electron microscopy imaging. E.G. designed and supervised the construction of mutants. S.J. and J.B. generated mutant and complementation strains. D.M.K. contributed to writing the original draft and the review editing, analyzed data, contributed to figure preparation, and provided supervision and conceptual input. R.C. contributed to conceptualization, writing the original draft and the review editing, supervised the research, and was responsible for funding acquisition.

## Acknowledgments

This research was financially supported by the National Science Center, Poland (Narodowe Centrum Nauki, Polska) via a research grant SONATA BIS 10 (2020/38/E/NZ9/00007) to Robert Czajkowski and by the University of Gdansk via a research grant UGrants-start (533-BGB0-GS04-25) to Marcin Borowicz. We thank Jan Jezierski (University of Gdansk, Poland) for his technical assistance.

## Supplementary Information

### Supplementary Tables

**Supplementary Table S1.** Plasmids used in this study.

| Plasmids | Description | Source |
| --- | --- | --- |
| pRE112 | Suicide vector for allelic exchange, Cm<sup>R</sup>, *sacB*, *oriT*<br>RP4, *oriR6K* | [27] |
| pEGL332 | Amp<sup>R</sup>, pSC101 ori, lacZp expression vector, mobilizable | [28] |
| pSJG | pEGL332-ΔDda3937_03810-ΔDda3937_03811, Amp<sup>R</sup>; | This study |

**Supplementary Table S2.**
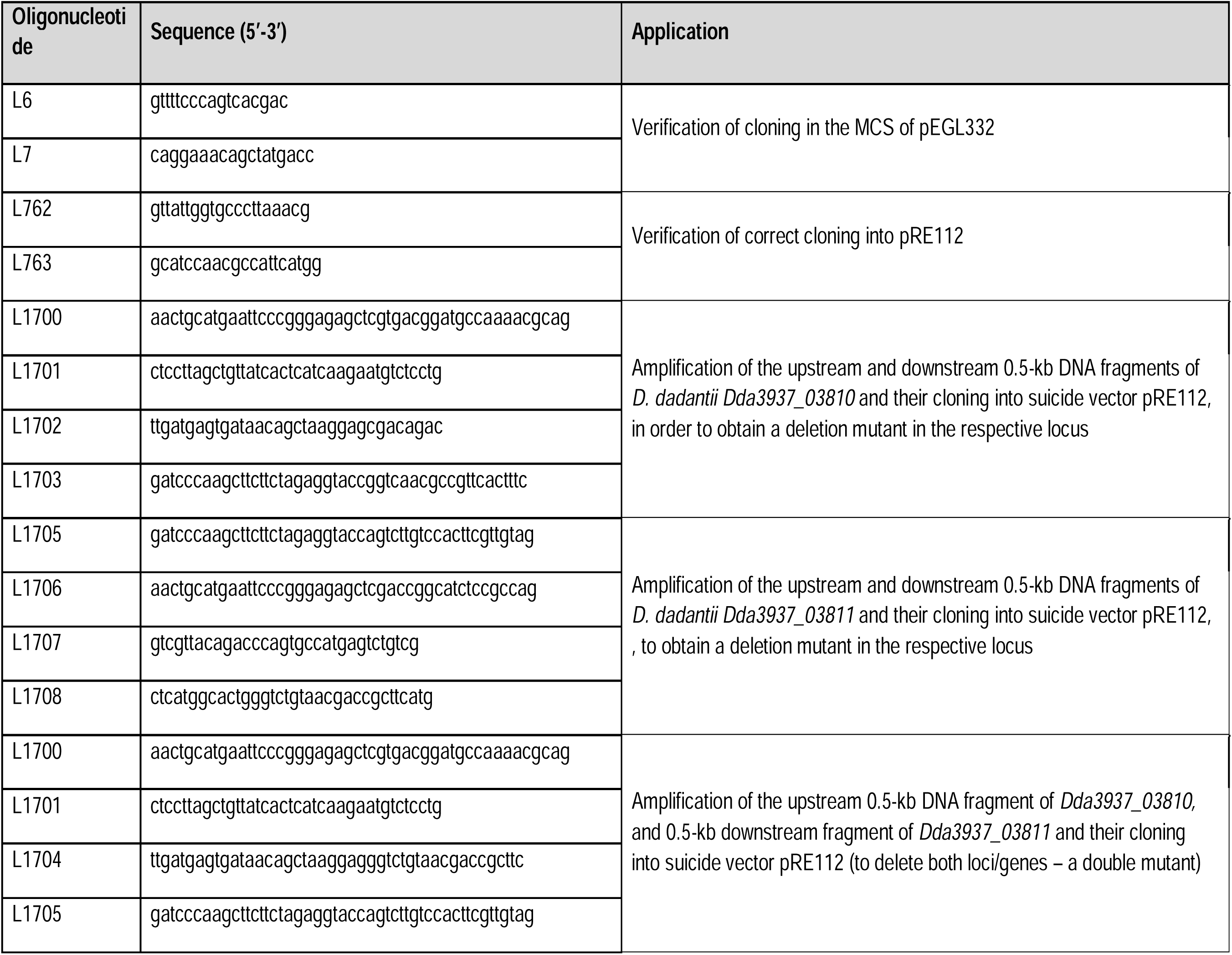
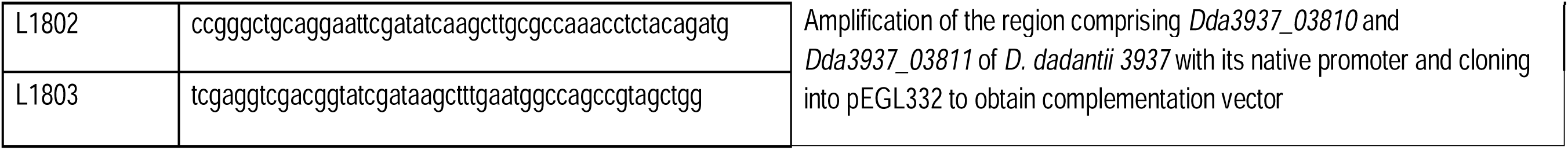
Oligonucleotides used in this study.

### Supplementary Figures

**Supplementary Fig. S1.**
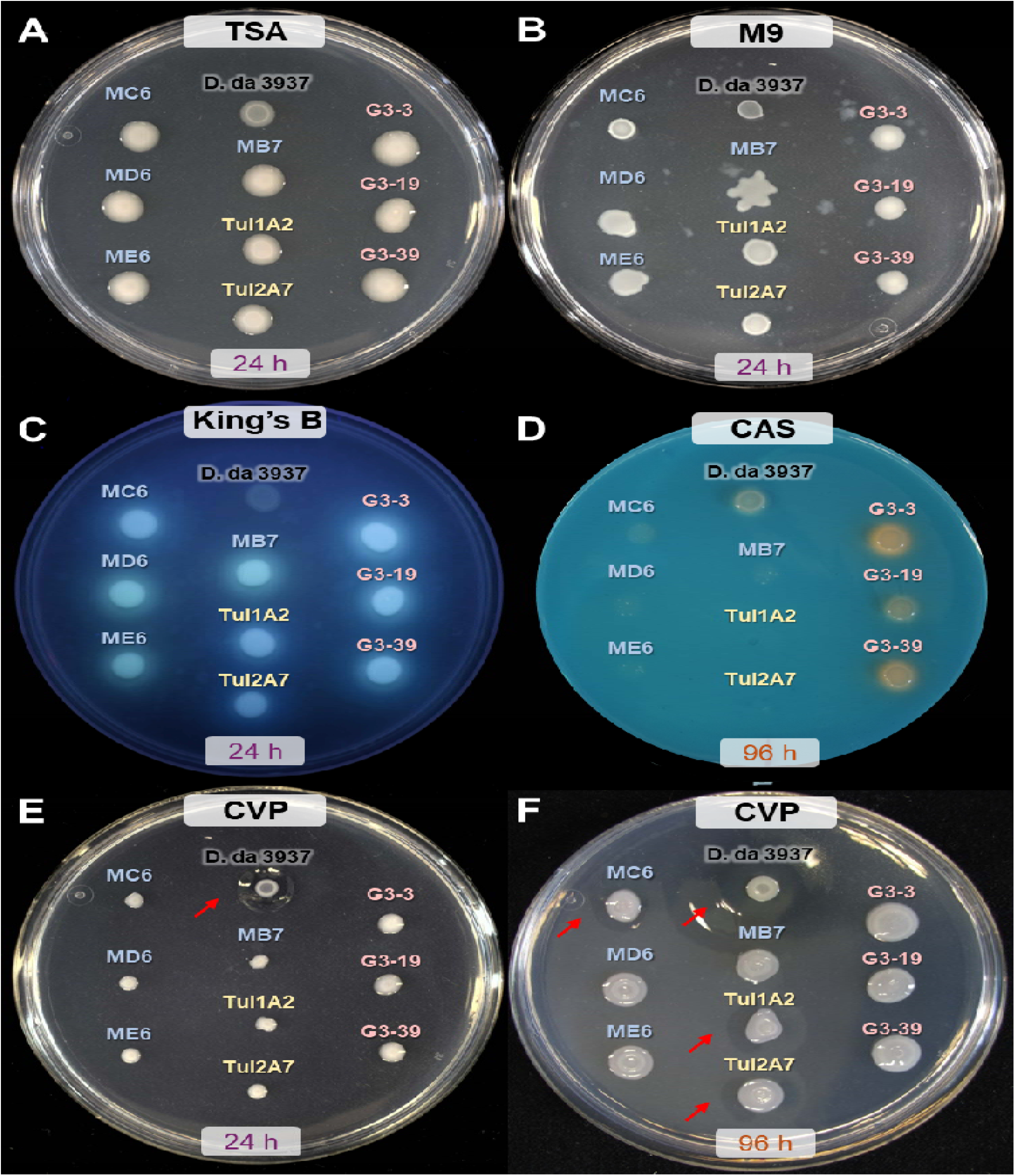
Morphology of P2D1 tailocin-susceptible *Pseudomonas* isolates grown on different media plates: TSA (A), M9 0.4% glucose (B), King’s B (C), CAS agar for siderophore production (D), and crystal violet pectate (CVP) for the production of pectinases (E and F). **I**ncubation time is indicated under each plate. On TSA, colonies of all tested *Pseudomonas* isolates exhibit a “slimy” morphology indicative of abundant extracellular matrix production. Images on King’s B medium were taken under UV light (λ = 365 nm) to visualize pyoverdine fluorescence, characteristic of fluorescent pseudomonads. Where present, cavities on CVP medium are marked by arrows. *D. dadantii* 3937 was included as a reference for colony morphology and as a negative control for King’s B and a positive control on CVP.

**Supplementary Fig. S2.**
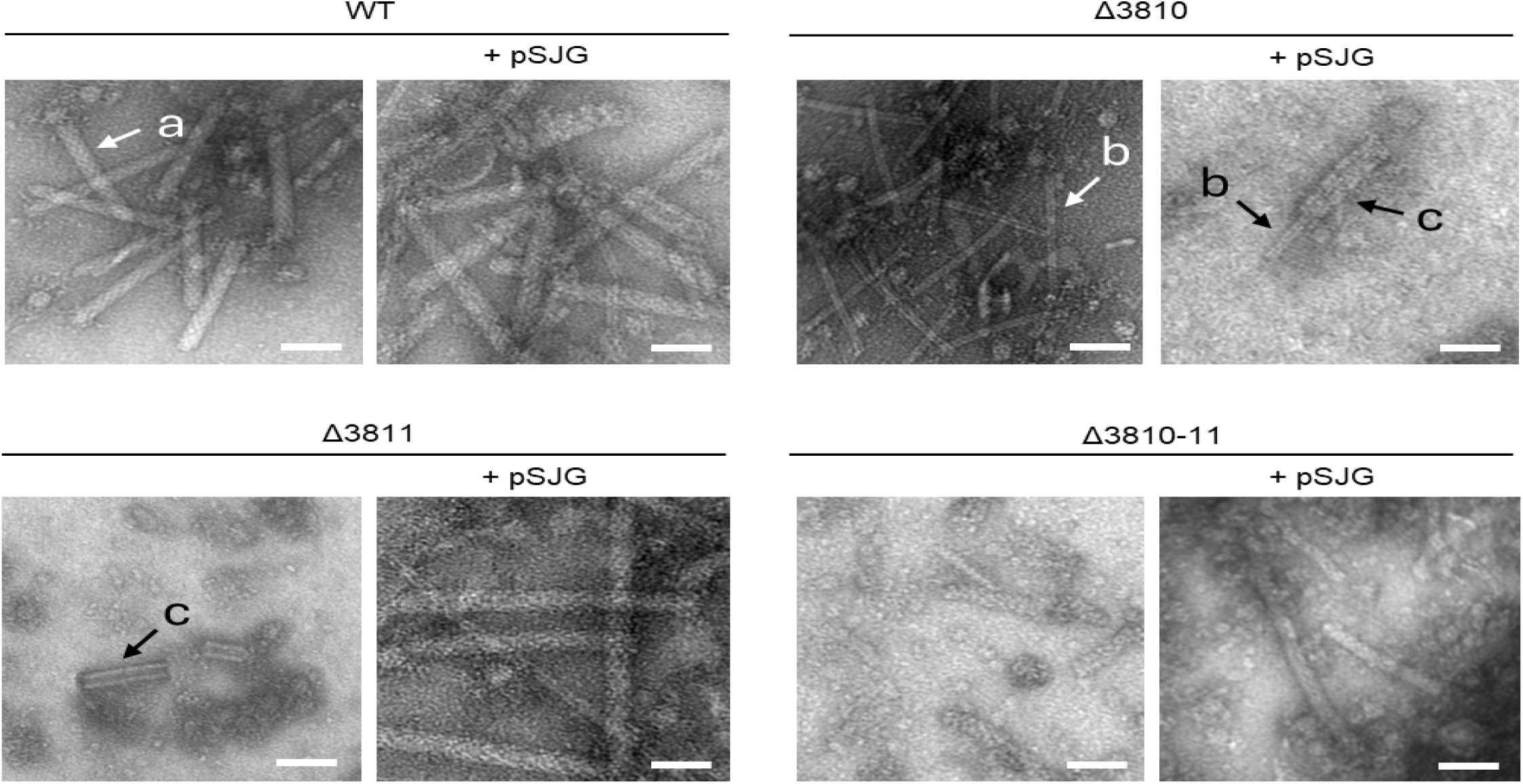
TEM images of tailocins purified from mitomycin C-induced cultures of the wild-type *D. dadantii* 3937 (WT) and its mutants. Only tubes are visible in the sheath mutant Δ3810, while only sheath fragments are present in the tube mutant Δ3811. No complete tailocin particles were detected in the preparations from the double mutant Δ3810-11. Scale bar 50 nm.

**Supplementary Fig. S3.**
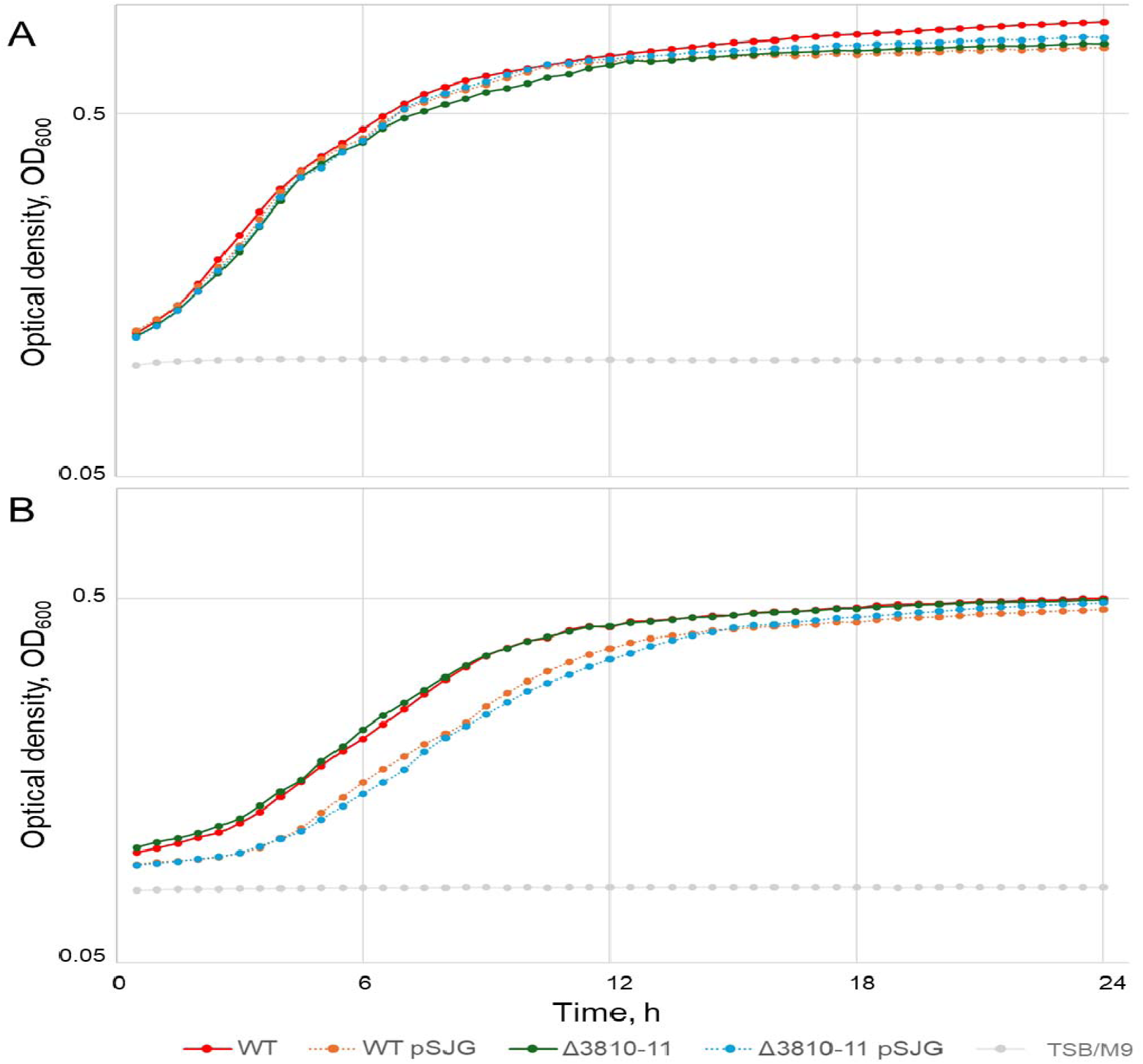
Comparison of growth curves for *D. dadantii* 3937 and its tailocin-defficient mutants. Overnight cultures were prepared in (A) tryptic soy broth (TSB, Oxoid), (B) M9 minimal medium (MP Biomedicals) supplemented with 0.4% glucose (Sigma-Aldrich) at 28 °C with shaking at 120 rpm. These cultures were diluted in a 96-well plate by mixing 10 µL of the overnight culture with 190 µL of fresh TSB. Each strain was tested in 12 technical replicates, and the experiment was performed in triplicate. The plates were incubated at 28 °C with continuous shaking at 237 rpm in an Epoch 2 microplate reader (BioTek). Optical density (OD) at 600 nm was measured every 30 minutes over 24 hours. Results are shown as average. pSJG – complementation plasmid. TSB/M9 – control with sterile medium.

**Supplementary Fig. S4.**
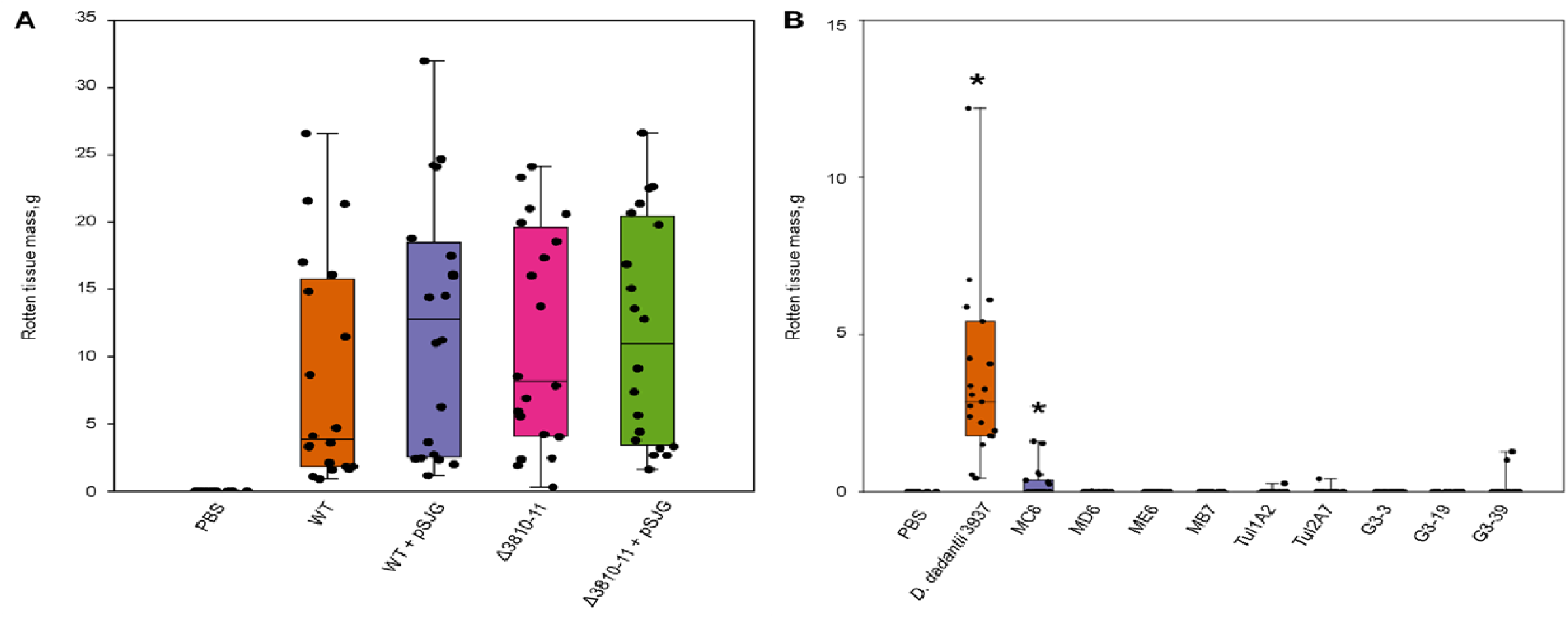
Virulence on potato tubers in a whole-tuber injection assay. (A) Virulence of *D. dadantii* 3937 and its tailocin-deficient double mutant Δ3810-11; pSJG denotes the complementation plasmid. No significant differences were detected between the wild type and its derivatives (Mann–Whitney test, p > 0.05). (B) Virulence of P2D1 tailocin-susceptible *Pseudomonas* isolates. Significant differences were observed between the negative control (PBS) and *D. dadantii* 3937, as well as isolate MC6 (one-sample Wilcoxon test, p < 0.05; marked with asterisks). Data are shown as box plots: whiskers indicate minimum and maximum values, boxes the interquartile range, and horizontal lines the medians. Each point represents an individual potato tuber (n = 20).

**Supplementary Fig. S5.**
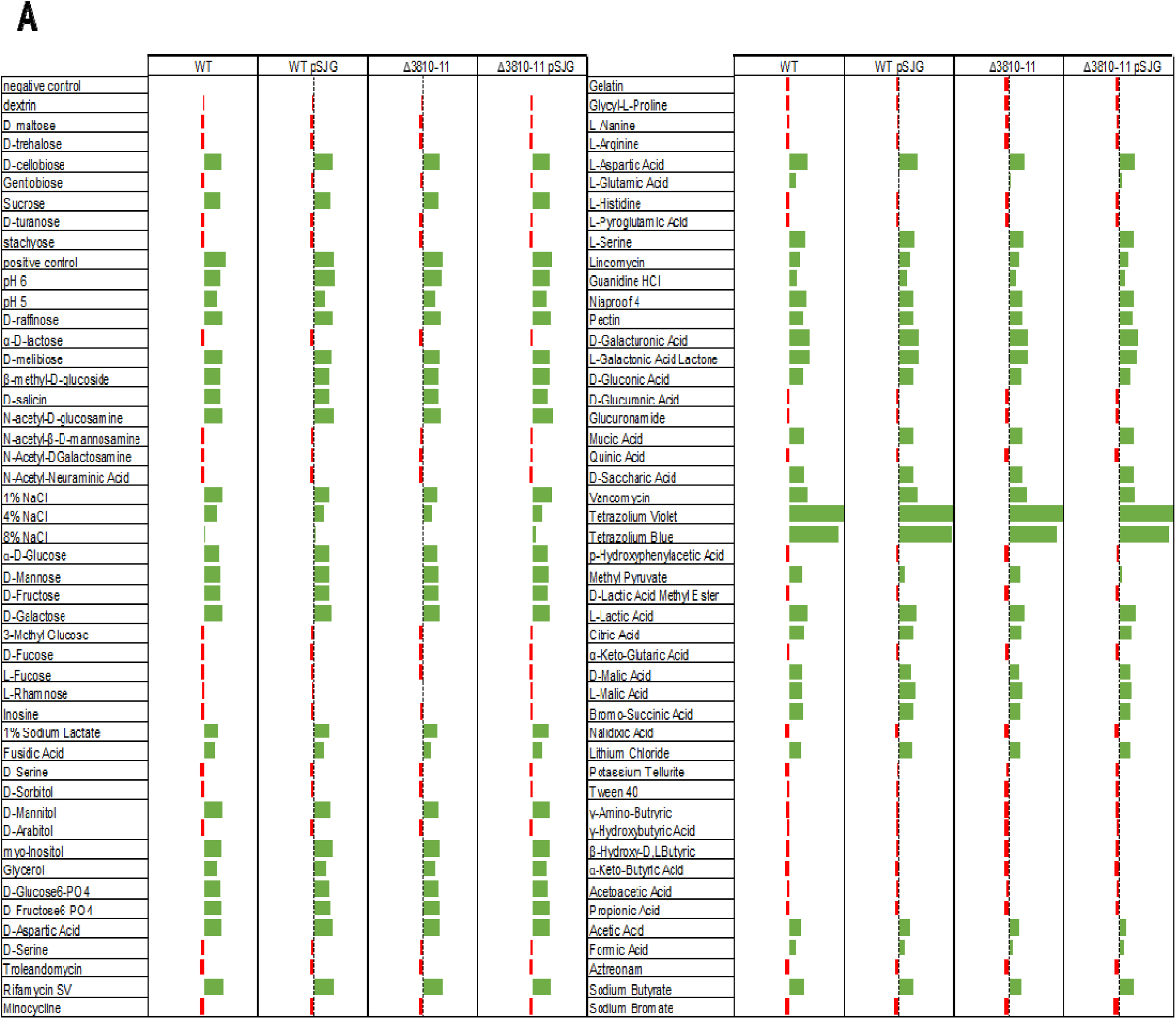
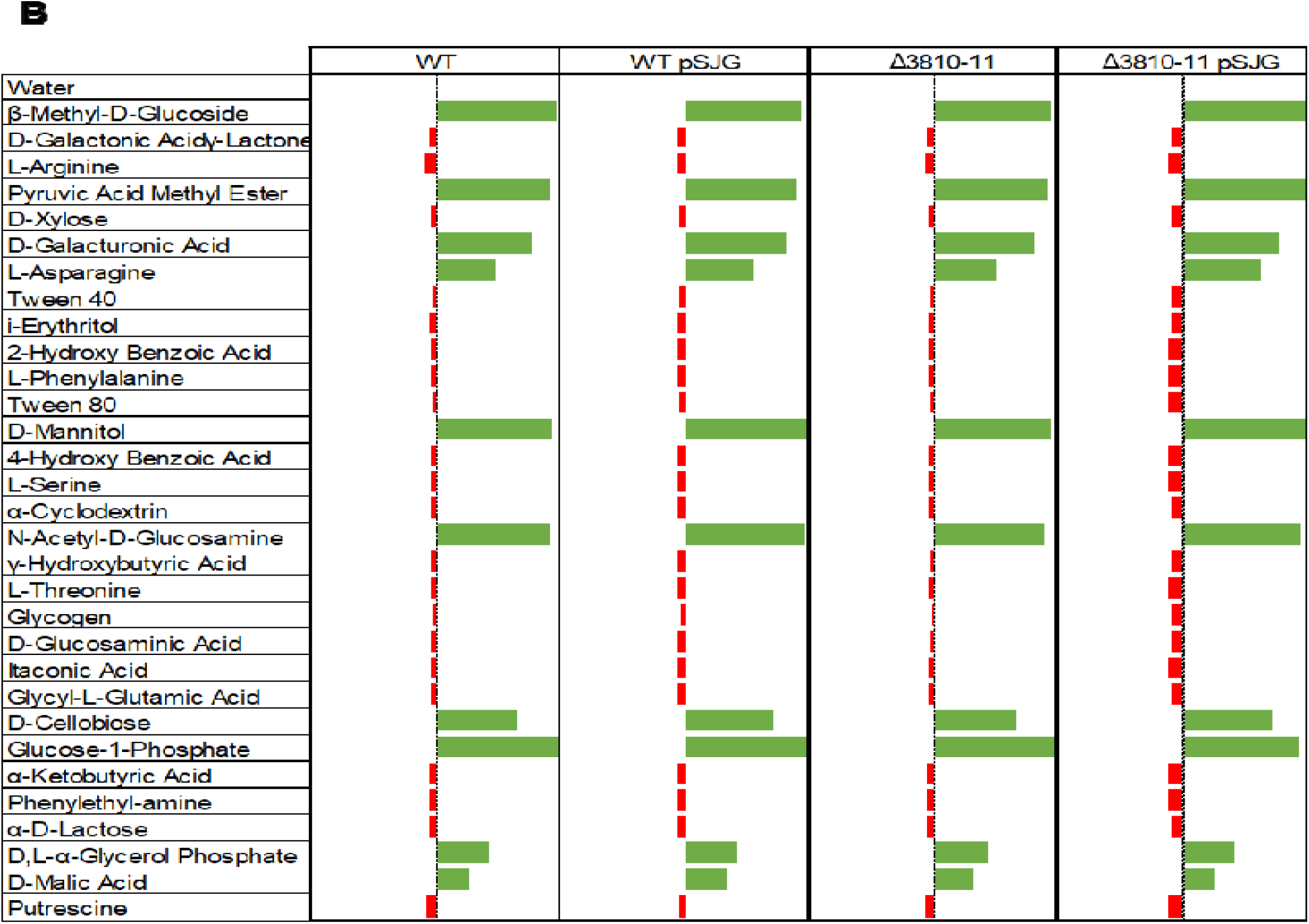
Phenotypic profiling of *D. dadantii* 3937 and its tailocin-deficient mutant using BIOLOG assays. (A) GEN III MicroPlate™ profiling (94 traits, including carbon utilization, chemical sensitivity, and physiological properties). (B) EcoPlate™ profiling (31 carbon-source utilization traits). Results were normalized to the negative control (positive reaction ≥2 × control) and averaged across three biological replicates. Data are presented as bar plots relative to the negative control baseline. pSJG denotes the complementation plasmid.

**Supplementary Fig. S6.**
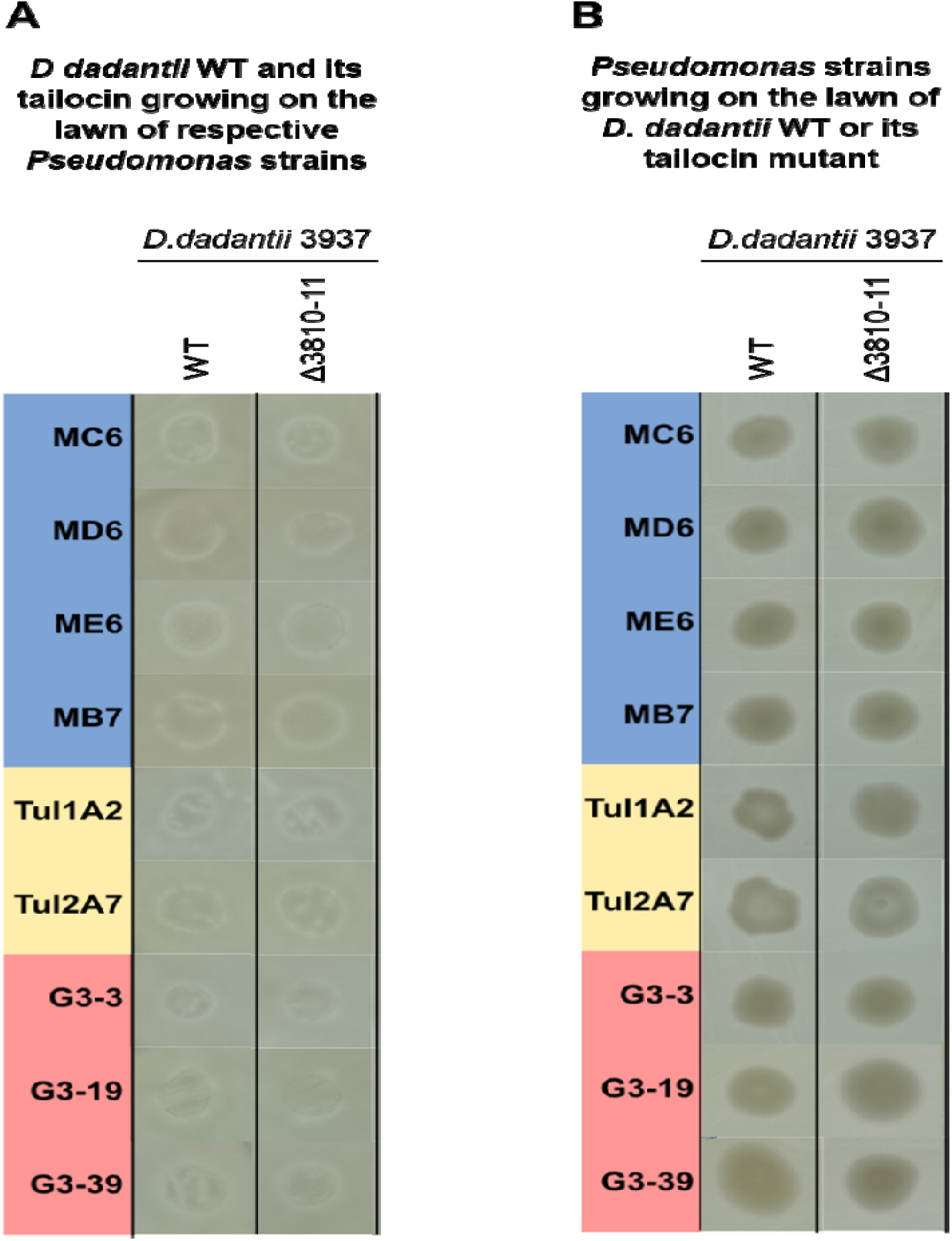
Reciprocal antibiosis assay on TSA media plates between *D. dadantii*, its tailocin-deficient mutant, and environmental *Pseudomonas* strains. *D. dadantii* wild-type and the mutant were spotted onto the lawns of environmental *Pseudomonas* strains (A) and *vice versa* (B). Results from one of two independent experiments yielding identical outcomes are shown.

### Supplementary Datasets

Supplementary Dataset 1 Isolates Taxonomical Placement Among *Pseudomonas* Type Strains Based On 16S rRNA Gene – Full Tree (pdf file).

Supplementary Dataset 2 Average Nucleotide Identity Calculation (Excel file)

Supplementary Dataset 3 Isolates Full Phenotype Profiles (Excel file)

Supplementary Script 1 Hierarchical clustering and dendrogram visualization (txt file)

