## SupplementaryDataSet1 for "Beyond kin killing: *Dickeya*-derived phage-tail-like bacteriocin P2D1 targets phylogenetically distant *Pseudomonas* spp"

**Phylogeny based on 16S rRNA gene**

To determine the taxonomic placement of environmental isolates within the *Pseudomonas* genus, we performed phylogenetic analysis based on 16S rRNA gene sequences. A total of 371 reference sequences representing Pseudomonas type strains were retrieved from the LPSN database (https://lpsn.dsmz.de/genus/pseudomonas; accessed 18 April 2025). The 16S rRNA sequences of the isolates and reference strains were aligned using the MUSCLE algorithm implemented in MEGA 12 (Kumar 2024).

The phylogeny was inferred using the Maximum Likelihood method and General Time Reversible model (Nei 2000) of nucleotide substitutions and the tree with the highest log likelihood (-18 605.00) is shown. The percentage of replicate trees in which the associated taxa clustered together, where the number of replicates (128) was determined adaptively (Kumar 2024), is shown next to the branches. The initial tree for the heuristic search was selected by choosing the tree with the superior log-likelihood between a Neighbor-Joining (NJ) tree (Saitou 1987) and a Maximum Parsimony (MP) tree. The NJ tree was generated using a matrix of pairwise distances computed using the General Time Reversible model (Nei 2000). The MP tree had the shortest length among 10 MP tree searches, each performed with a randomly generated starting tree. The evolutionary rate differences among sites were modeled using a discrete Gamma distribution across 5 categories (*+G*, parameter = 0.2619), with 50.08% of sites deemed evolutionarily invariant (*+I*). The analytical procedure encompassed 380 nucleotide sequences. The partial deletion option was applied to eliminate all positions with less than 95% site coverage resulting in a final data set comprising 1 286 positions. Evolutionary analyses were conducted in MEGA12 (Kumar 2024) utilizing up to 12 parallel computing threads.

1. Nei M. and Kumar S. (2000). Molecular Evolution and Phylogenetics. Oxford University Press, New York.

2. Kumar S., Stecher G., Suleski M., Sanderford M., Sharma S., and Tamura K. (2024). Molecular Evolutionary Genetics Analysis Version 12 for adaptive and green computing. Molecular Biology and Evolution 41:1-9.

3. Saitou N. and Nei M. (1987). The neighbor-joining method: A new method for reconstructing phylogenetic trees. Molecular Biology and Evolution 4:406-425.

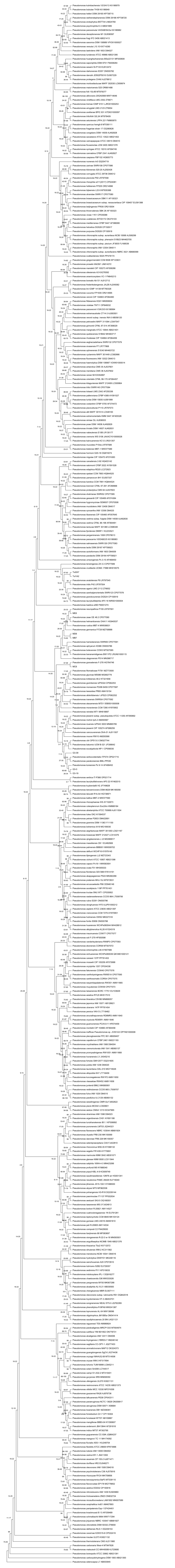
